# Dynamics of annatto pigment synthesis and accumulation revealed by integrated chemical, anatomical, and RNA-Seq analyses

**DOI:** 10.1101/2022.09.14.507995

**Authors:** Viviane Santos Moreira, Vinicius Carius de Souza, Virgínia Lúcia Fontes Soares, Aurizangela Oliveira Sousa, Katiucia Ticila de Souza de Nascimento, Monique Reis de Santana, Tiyoko Nair Hojo Rebouças, Carlos André Espolador Leitão, Priscila Vanessa Zabala Capriles Goliatt, Daniele Vidal Faria, Wagner Campos Otoni, Marcio Gilberto Cardoso Costa

**Affiliations:** Instituto Federal de Educação Ciência e Tecnologia da Bahia, Euclides da Cunha, Bahia 48500-000, Brazil; Departamento de Ciências da Computação, Universidade Federal de Juiz de Fora, Juiz de Fora, Minas Gerais 36036-900, Brazil; Centro de Biotecnologia e Genética, Departamento de Ciências Biológicas, Universidade Estadual de Santa Cruz, Ilhéus, Bahia 45662-900, Brazil; Centro Multidisciplinar do Campus Luís Eduardo Magalhães, Universidade Federal do Oeste da Bahia, Luís Eduardo Magalhães, Bahia 47850□000, Brazil; Departmento de Fitotecnia e Zootecnia, Universidade Estadual do Sudoeste da Bahia, Vitória da Conquista, Bahia 45083-900, Brazil; Departamento de Ciências Naturais, Universidade Estadual do Sudoeste da Bahia, Vitória da Conquista, Bahia 45083-900, Brazil; Departamento de Biologia Vegetal/BIOAGRO, Universidade Federal de Viçosa, Viçosa, Minas Gerais 36570-900, Brazil

**Keywords:** apocarotenoids, *Bixa orellana*, bixin, carotenoid, transcriptome

## Abstract

Bixin is a commercially valuable apocarotenoid pigment found in the seed aril of *Bixa orellana*. The dynamics and regulation of its biosynthesis and accumulation during seed development remain largely unknown. Here, we combined chemical, anatomical, and transcriptomic data to provide stage-specific resolution of the cellular and molecular events occurring during *B. orellana* seed development. Seeds at five developmental stages (S1–S5) were used for analysis of bixin content and seed anatomy, and three of them (S1, S3 and S4) selected for Illumina HiSeq sequencing. Bixin accumulated sharply during seed development, particularly during the S2 stage, peaking at the S4 stage, and then decreasing slightly in the S5 stage. Anatomical analysis revealed that bixin accumulated in the large central vacuole of specialized cells, which were scattered throughout the developing mesotesta at the S2 stage, but enlarged progressively at later stages, until they occupied most of the parenchyma in the aril. A total of 13 million reads were generated and assembled into 73,381 protein-encoding contigs, from which 312 were identified as containing 1-deoxy-D-xylulose-5-phosphate/2-C-methyl-D-erythritol-4-phosphate (DOXP/MEP), carotenoid, and bixin pathways genes. Differential expression analysis of these genes revealed that 50 of them were differentially expressed between all the seed developmental stages, including seven carotenoid cleavage dioxygenases, eight aldehyde dehydrogenases and 22 methyltransferases. Taken together, these results provide a comprehensive description of the cellular and molecular events related to the dynamics of bixin synthesis and accumulation during seed development in *B. orellana*.

## Introduction

*Bixa orellana*, commonly known as annatto, is a perennial tree native to tropical America. It represents the only seed-specific source of a red-yellow dye also referred to as annatto, which has been widely used since pre-Columbian times. Nowadays, annatto is employed as an additive in several industries, ranking second in economic importance among natural color additives and remains comparatively inexpensive (Mercadante and Pfander 1998; Giuliano et al. 2003). The major (>80%) color component of annatto pigment is bixin, a dicarboxylic monomethyl ester apocarotenoid, followed by traces of other carotenoids, including norbixin, bixin dimethyl ester, and several apocarotenoids (Mercadante and Pfander 1998).

Annatto pigment is stored in the aril, which may account for up to 10% of the dry weight of the seed (Carvalho et al. 2010). Bixin occupies the entire volume of mature aril cells, suggesting that they store the pigment but lose their biosynthetic function (Rodríguez-Ávila et al. 2011). Unlike most plant carotenoid pigments, which are synthesized and stored as plastoglobules within chromoplasts or as crystalline, fibrous or membranous structures (Vishnevetsky et al. 1999), *B. orellana* carotenoids are formed in chromoplasts, but are stored in the vacuole (Bouvier et al. 2003b; Giuliano et al. 2003; Louro and Santiago 2016). This uncommon feature has been reported only in arbuscular mycorrhizae of roots (Fester et al. 2002) and saffron (*Crocus*) pistils (Bouvier et al. 2003b). Carotenoids are visible already at the globular embryo stage as electron-dense plastoglobules in chromoplasts, as well as cytoplasmic electron-dense lipid bodies and electron-lucent vesicles of specialized aril cells called carotenoid storage cells (CSCs) (Louro and Santiago 2016). During aril development, the number and volume of CSCs increase, as their vacuoles are progressively filled with the above globules and vesicles, and this trend appears to be associated with an increase in carotenoid concentration (Louro and Santiago 2016).

Knowledge about the bixin biosynthetic pathway and its regulatory mechanisms remains scarce. Based on its structural similarity to the saffron pigment crocetin, it has been suggested that bixin biosynthesis could involve a carotenoid cleavage dioxygenase (CCD), an aldehyde dehydrogenase (ALDH), and a methyltransferase (MT), catalyzing sequential steps from an initial C_40_ carotenoid such as lycopene (Mercadante et al. 1997; Jako et al. 2002; Bouvier et al. 2003a). This assumption is supported by trace amounts of several lycopene cleavage derivatives in annatto (Mercadante et al. 1997; Mercadante and Pfander 1998), as well as by the accumulation of bixin in an *Escherichia coli* strain engineered for the production of lycopene and heterologous co-expression of *B. orellana* lycopene cleavage dioxygenase, bixin ALDH, and norbixin MT (Bouvier et al. 2003a).

In plants, carotenoids are synthesized in plastids via the DOXP/MEP pathway, which involves the action of several enzymes, including 1-deoxy-D-xylulose-5-phosphate synthase (DXS), 1-deoxy-D-xylulose-5-phosphate reductoisomerase (DXR), 2-C-methyl-D-erythritol 4-phosphate cytidyltransferase (MCT), 2-C-methyl-D-erythritol 2,4-cyclodiphosphate synthase (MDS), 4-hydroxy-3-methylbut-2-enyl diphosphate reductase (HDR), 4-hydroxy-3-methylbut-2-en-1-yl diphosphate synthase (HDS), and isopentenyl diphosphate isomerase (IDI) (Lichtenthaler 1999). The main products of the DOXP/MEP pathway, isopentenyl pyrophosphate (IPP) and dimethylallyl pyrophosphate (DMAPP), are condensed into the C_20_ compound geranylgeranyl pyrophosphate (GGPP), the immediate precursor of all carotenoids, by the action of GGPP synthase (Lichtenthaler 1999). The condensation of two C_20_ units of GGPP by phytoene synthase (PSY) produces phytoene, the first C_40_ carbon intermediate of the carotenoid biosynthetic pathway (Sun et al. 2018). Phytoene is then converted into lycopene by poly-*cis* desaturation/isomerization reactions relying on two desaturases, phytoene desaturase (PDS) and zeta-carotene desaturase (ZDS), and two isomerases, carotene *cis-trans* isomerase (CRTISO) and 15-*cis*-zeta-carotene isomerase (Z-ISO) (Sun et al. 2018).

Lycopene is the substrate of two competing cyclases, lycopene ε-cyclase (LCY-ε) and lycopene β-cyclase (LCY-β), which introduce, respectively, ε-rings or β-rings at the end of the molecule. A series of reactions catalyzed by heme and non-heme hydroxylases, an epoxidase, a de-epoxidase, and a neoxanthin synthase complete the canonical plant carotenoid biosynthetic pathway. Carotenes and xanthophylls are subjected to further modifications, including non-enzymatic or enzymatic oxidative cleavage by CCDs to produce apocarotenoids, which may act as phytohormones, signaling molecules, volatiles, and pigments (Sun et al. 2018).

Transcriptome sequencing of *B. orellana* seeds has led to the identification of several candidate genes involved in the DOXP/MEP, carotenoid, and bixin biosynthetic pathways (Jako et al. 2002; Soares et al. 2011; Cárdenas-Conejo et al. 2015). EST analysis of 870 clones from a cDNA library derived from *B. orellana* seeds identified six contigs encoding enzymes belonging to the DOXP/MEP pathway (DXS, HDR, two DXRs, and two HDSs), two relating to GGPP synthase, six belonging to the carotenoid pathway (PSY, three PDSs, and two ZDSs), and 12 belonging to the bixin pathway (five CCDs, five ALDHs, and two carboxyl MTs) (Jako et al. 2002). In another study, EST analysis of 792 clones of a non-normalized cDNA library from *B. orellana* seeds revealed two candidate genes for the bixin pathway: CCD subclass 4 (*BoCCD4*) and caffeic acid 3-O-methyltransferase (*BoCOMT*) (Soares et al. 2011). Quantitative real-time PCR (qRT-PCR) analysis showed that both genes were upregulated during seed development, and coincided with growing maturity and seed pigmentation (Soares et al. 2011). More recently, high-throughput RNA sequencing (RNA-Seq) from young leaves, as well as immature and mature seeds of *B. orellana*, yielded 52,549 contigs and identified 11 candidate genes involved in the DOXP/MEP pathway (four DXSs, DXR, MCT, CMK, MDS, HDS, HDR, and IDI), 22 in the carotenoid pathway [two PSYs, two PDSs, ZDS, 15-*cis*-zeta-carotene isomerase, three carotene *cis-trans* isomerases, two LCY-εs, two LCY-βs, CHY1-β hydroxylase, three CYPs, two zeaxanthin epoxidases (ZEPs), two violaxanthin de-epoxydases (VDEs), and NSY], and 41 in the bixin pathway (nine CCDs, 20 ALDHs, and 12 SABATH MTs) (Cárdenas-Conejo et al. 2015). Of the 22 candidate genes selected for qRT-PCR analysis using new RNA samples from the same tissues, 13 (*BoDXS2a, BoPDS1, BoZDS, BoCCD1-3, BoCCD1-4, BoCCD4-1, BoCCD4*-*2, BoCCD4*-*3, BoALDH3H1-1, BoALDH3I1, BoSABATH-1, BoSABATH-3*, and *BoSABATH-4*) were preferentially expressed in immature seeds, suggesting their involvement in the DOXP/MEP, carotenoid, and bixin pathways, respectively (Cárdenas-Conejo et al. 2015). Surprisingly, transcripts corresponding to the cDNA sequences previously identified and functionally characterized by Bouvier et al. (2003a) were not found in any transcriptome studies of *B. orellana* (Jako et al. 2002; Soares et al. 2011; Cárdenas-Conejo et al. 2015), suggesting that their cDNA library could be contaminated with exogenous mRNA.

Bixin synthesis and accumulation in the seeds of *B. orellana* remains a poorly explored process. Previous studies have mostly examined dissociated and static snapshots of chemical, cellular, or molecular data. In the present study, we integrated chemical, anatomical, and transcriptomic data obtained during different stages of *B. orellana* seed development to provide a more comprehensive view of the cellular and molecular events that contribute to bixin production and accumulation.

## Materials and methods

### Plant material

Samples of mature leaves, flower buds, flowers, and seeds at five developmental stages (S1– S5) from three biological replicates (individual plants) of *B. orellana* L. var. Embrapa 37 were harvested as previously described (Moreira et al. 2018). S1, S2, S3, S4 and S5 seeds were collected from capsules between 7 and 14, 14 and 28, 28 and 42, 42 and 63, and 63 and 77 days after anthesis (DAA), respectively. S1 to S4 capsules represent the immature fruit stage, and S5 the mature fruit stage. The collected samples were either immediately submerged in liquid nitrogen and maintained at -80 °C for subsequent isolation of total RNA, instantly fixed in Karnovsky’s solution (Karnovsky 1965) for anatomical analysis, or maintained *in natura* for chemical analysis of bixin content.

### Analysis of bixin content

The samples were dried in a closed-circulation oven for 24 h at 55 °C. Bixin content was determined using the 5% KOH method (Yabiku and Takahashi 1991). Briefly, 25 g of each sample (dry weight basis) was weighed and submerged in 150 mL of a 5% KOH boiling solution for 1 min. Without stirring, the samples were cooled under running tap water, filtered, and the mass retained on the filter was washed with distilled water (100 mL) seven to nine times. The volume was then calibrated to 1000 mL, 2 mL was transferred to a 1000-mL volumetric flask, and the volume was topped up with 0.5% KOH. Readings were performed in a spectrophotometer (SP-220; Biospectro, Curitiba, PR, Brazil) at 453 nm in a 1-cm quartz cuvette. The 0.5% KOH solvent was used as reference. The total pigment content expressed as bixin was calculated using the following formula:

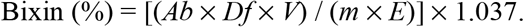

where *Ab* is the absorbance, *Df* is the dilution factor, *V* is the initial extract volume, *m* is the seed mass, *E* is the absorption coefficient (3473) for bixin in KOH, and 1.037 is the norbixin-to-bixin conversion factor based on molecular weight (380.48 and 394.51, respectively).

### Anatomical analysis

Samples were fixed, dehydrated using an ethanol series, embedded in glycol-methacrylate resin (Historesin; Leica, Heidelberg, Germany), and sectioned in a rotating microtome to obtain serial cuts of 3-μm thickness. Sample sections were stretched on microscope slides and stained with toluidine blue in McIlvaine buffer (pH 4) for structural characterization and histochemical inferences by orthochromasia and metachromasia. The slides were photographed using a DM750 photomicroscope (Leica Microsystems, Heerbrugg, Switzerland) equipped with a digital image capture system. Three slides per organ per plant from a total of three plants were used as replicates.

### RNA isolation, sequencing, and data analysis

Total RNA was isolated as previously described (Moreira et al. 2018), using the RNAqueous® kit Plant RNA Isolation Aid reagent (Ambion Inc., Austin, TX, USA). RNA-Seq libraries were constructed using the Truseq™ Stranded mRNA LT Sample Prep Kit (Illumina, San Diego, CA, USA), following the manufacturer’s instructions. Libraries were quantified on a Stratagene Mx3005P qRT-PCR system (Agilent Technologies, Santa Clara, CA, USA) using the KAPA Library Quantification Kit (Kapa Biosystems, Wilmington, MA, USA). The average library fragment size of the qPCR products (∼400 bp) was assessed by electrophoresis on a 3% agarose gel. Sequencing was carried out by GenOne Biotechnologies, Rio de Janeiro, RJ, Brazil, using an Illumina HiSeq 2500 platform with 150-bp pair-end reads mode. Libraries from two independent biological replicates (plants) at the S1, S3, and S4 stages were sequenced.

The quality of each sequenced library was evaluated using FastQC software (Andrews, 2010). Low-quality bases and sequencing adapters were removed using Trimmomatic software (Bolger et al. 2014). *De novo* assembly was carried out with clean reads from the combined RNA-Seq datasets using Trinity v2 (Grabherr et al. 2011) with default parameters. Coding regions in the assembled transcripts were predicted using the Transdecoder program from the Trinity package. After ORF identification and translation to the amino acid sequence, the non-redundant (NR) database was queried using BlastP software to identify homologous sequences with an e-value cutoff of 1e-6. Gene Ontology (GO) annotation and Kyoto Encyclopedia of Genes and Genomes (KEGG) metabolic pathways were obtained using Blast2GO software by applying default parameters. Genes potentially involved in the DOXP/MEP, carotenoid, and bixin pathways became ‘candidate genes’ for further differential expression analysis.

Transcript abundance estimation was carried out using a salmon algorithm that generated a count matrix containing RNA-Seq fragment counts for each transcript by a quasi-mapping approach. The transcript count matrix was used for differential expression analysis based on the limma/voom method in R. Accordingly, a pairwise comparison between all the libraries (S1_r1, S1_r2, S3_r1, S3_r2, S4_r1, and S4_2) was performed. Differentially expressed genes (DEGs) were extracted, based on a significant false discovery rate (FDR) < 0.05. Heatmaps and Venn diagrams were generated in R. The subcellular localization of proteins encoded by DEGs was predicted using the Plant-mSubP tool (http://bioinfo.usu.edu/Plant-mSubP/) (Sahu et al. 2020).

### qRT-PCR analysis

qRT-PCR analyses were performed as previously described (Moreira et al. 2018). All the qRT-PCR reactions were run on a Stratagene Mx 3005P apparatus (Agilent Technologies, Santa Clara, CA, USA) using the 5x HOT FIREPol® EvaGreen® qPCR Supermix kit (Solis BioDyne, Tartu, Estonia), according to the manufacturer’s instructions. The qRT-PCR reactions were carried out in a reaction volume of 20 μL, containing 4 μL SYBR green, 1 μL (5 mM) each primer, 5 μL sterile Milli-Q water and 10 μL (10 ng) cDNA sample. The amplification conditions were performed using the following steps: activation of Taq DNA Polymerase at 95ºC for 12 minutes, followed by 40 cycles of denaturation at 95ºC for 15 seconds, annealing at 60ºC for 30 seconds and extension at 72ºC for 30 seconds. The non-template control (NTC) reactions were included in the analysis in triplicate. Primers designed to avoid the conserved regions were generated using Primer3 version 0.4.0 (http://bioinfo.ut.ee/primer3-0.4.0/). *RPL38*, encoding the 60S ribosomal protein L38, was used as an endogenous gene for normalization of qRT-PCR data (Moreira et al. 2018). The primers used in qRT-PCR reactions are listed in Table S1. The comparative cycle threshold (Ct; 2^−ΔΔCt^) method was used for expression quantification, considering the mean of the Ct values of the three technical replicates from two biological replicates.

### Statistical analysis

Statistical differences in bixin content and qRT-PCR results were assessed based on ANOVA and means separation by Student’s *t* tests, with a statistically significant value of *P* ≤ 0.05.

## Results

### Bixin content in different organs of *B. orellana*

Bixin quantification in samples of mature leaves, flower buds, flowers, and seeds at five developmental stages (S1–S5) showed that the pigment was present in all analyzed organs, albeit at very different levels (Fig. 1). Bixin content was relatively low in mature leaves (0.6%), flower buds (0.3%), flowers (0.4%), and seeds at the first (S1) developmental stage (0.4%), but became significantly higher at the S2 stage (2.7%), increasing continuously at S3 (3.9%) and S4 stages (5.0%), and decreasing slightly at the mature (S5) stage (4.8%).

**Fig. 1.**
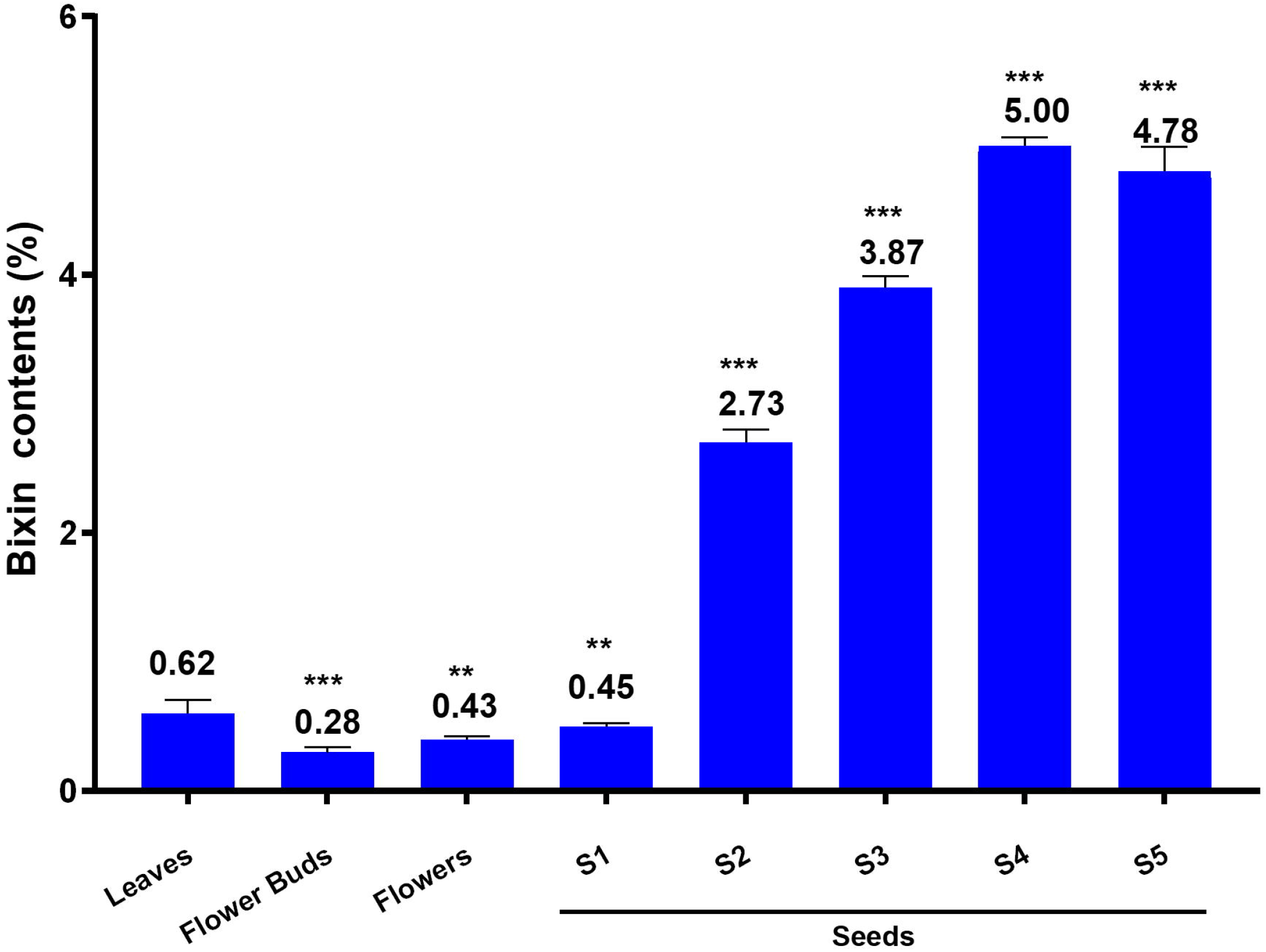
Bixin content (%) in different organs of *B. orellana* L. S1–S5: seed developmental stages from initial to mature, respectively. Statistically significant differences at *P* ≤ 0.01 (**) and *P* ≤ 0.001 (***) according to Student’s *t* tests.

### Anatomical analysis of seminal rudiment and seed development in *B. orellana*

Except for S1-stage seeds, whose samples were torn during tissue sectioning due to their very small size, we were able to perform anatomical analysis in representative sample sections from flower buds, flowers, and S2- to S5-stage seeds. In the seminal rudiment of flower buds, cells were larger and more vacuolated in the outer than in the inner integument (Fig. 2A). In the seminal rudiment of the flower, cells of the mesotesta showed increased vacuolization and became the most voluminous in the testa or aril (Fig. 2B). In S2-stage seeds, the mesotesta was characterized by the presence of numerous specialized CSCs of varying size, with cytoplasm densely stained by toluidine blue and various non-stained small and independent spherical vacuoles; the remaining area was occupied by small, ordinary parenchyma cells with a weakly colored cytoplasm (Fig. 2C). At the S3 stage, CSCs enlarged to occupy almost completely the region of the mesotesta, showing strong toluidine blue staining in the cytoplasm and a single, colorless, large central vacuole resulting from the fusion of small vacuoles (Fig. 2D). At this stage, ordinary parenchyma cells of the mesotesta were much smaller and compressed by the larger CSCs; whereas the cells in the exotegmen were elongated, assuming a columnar shape (Fig. 2D). Bixin became apparent at the S4 stage in CSCs of the mesotesta (Fig. 2E). These cells were much more expanded than at the previous stage, showing a peripheral and voluminous cytoplasm strongly stained with toluidine blue, plus a central vacuole fully filled with bixin, which under the microscope assumed a dark brown coloration (Fig. 2E). The small parenchyma cells of the mesotesta assumed a collapsed appearance and were crushed by CSCs (Fig. 2E). At the S5 stage, the fully developed mature seeds showed a marked accumulation of pigment in the vacuoles of CSCs (Fig. 2F). The latter showed a thinner cytoplasm and a collapsed appearance, while the macrosclereids that constituted the exotegmen were strongly sclerified (Fig. 2F).

**Fig. 2.**
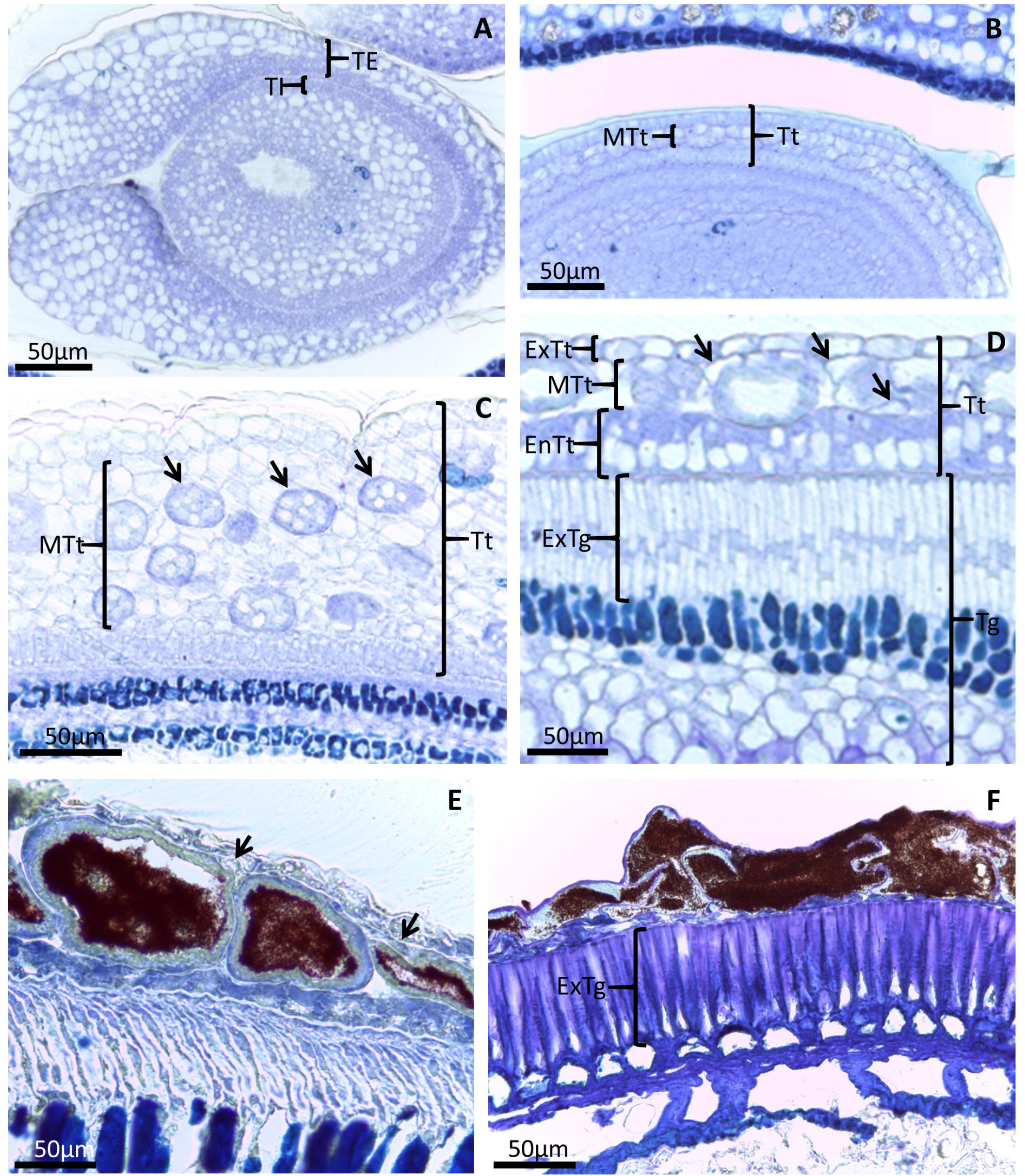
Anatomical sections of the seminal rudiment and seed of *B. orellana* at different developmental stages, stained with toluidine blue. (**A**) Seminal rudiment of the flower bud with the outer integument showing larger and more vacuolated cells than the inner integument. (**B**) Seminal rudiment of the flower displaying enlarged cells in the mesotesta, each containing a large vacuole. (**C**) Seed at the S2 stage showing developing mesotesta CSCs (arrows) with dense cytoplasm and numerous small vacuoles sparsely distributed throughout the cytoplasm. (**D**) Seed at the S3 stage showing CSCs with the newly formed large central vacuole and smaller, non-secretory mesotesta cells (arrows) surrounding the CSCs. (**E**) Seed at the S4 stage showing CSCs containing bixin in their large central vacuole and collapsed mesotesta non-secreting cells (arrows) adjacent to CSCs. (**F**) Fully developed seed at the mature S5 stage. EnTt, endotesta; ExTt, exotesta; ExTg, exotegmen; MTt, mesotesta; TE, external tegument; Tg, tegmen; TI, inner tegument; Tt, testa.

### RNA-Seq, de novo sequence assembly, and annotation

Six RNA libraries were sequenced for three stages of seed development (S1, S3, and S4), generating a total of 13,442,090 raw reads (Table S2). After filtering for low-quality reads, 7,572,594 high-quality (Q ≥ 20 or error < 0.1%) clean reads were retained (Table S2). Clean reads were assembled into 79,650 contigs with an average length of 994.91 bp and 50% of all bases coming from subreads longer than 1,603 bp (Table S3). From this dataset, the Transdecoder predicted a total of 73,381 ORFs with an average length of 715.06 bp and 50% of all bases coming from subreads longer than 1,191 bp (Table S3).

Functional annotation indicated that 51,632 ORFs (70.4%) were annotated against the NR database. BlastP analysis showed that 42% of *B. orellana* transcripts were similar to those of *Theobroma cacao*, 6% to those of *Corchorus* spp., and 3%–5% to those of *Gossypium* spp. (Fig. S1A), all of which belong to the Malvales order. All sequences with blast results were annotated against the InterPro database, and 42,816 sequences were annotated against the GO database (Fig. S1B).

### Transcriptome profiling of candidate *B. orellana* genes at different seed developmental stages

A total of 312 sequences encoding DOXP/MEP, carotenoid, and bixin pathway enzymes were identified among all annotated sequences and mapped into KEGG metabolic pathways (Fig. S2). These sequences were considered as candidate genes for differential expression analysis and deposited in the GenBank database under accession numbers MW885276 to MW885587. Prior to analysis of differential expression, 40 redundant sequences with 100% identity and 23 sequences whose counts per million were greater than zero in only one biological replicate were removed. Venn diagram analysis of the remaining 249 sequences showed that 226 (90.8%) were shared among all three seed stages (Fig. 3A); whereas five (2%) were shared by S1 and S3 stages, 13 (5.2%) were shared by S1 and S4 stages, and five (2%) were shared by S3 and S4 stages (Fig. 3A). Pearson correlation analysis of these candidate genes’ expression across all samples revealed consistency among the biological replicates and transcriptional differences between different seed stages (Fig. 3B). Generally, there was less transcriptional variation between the S3 and S4 stages than with respect to the S1 stage (Fig. 3B).

**Fig. 3.**
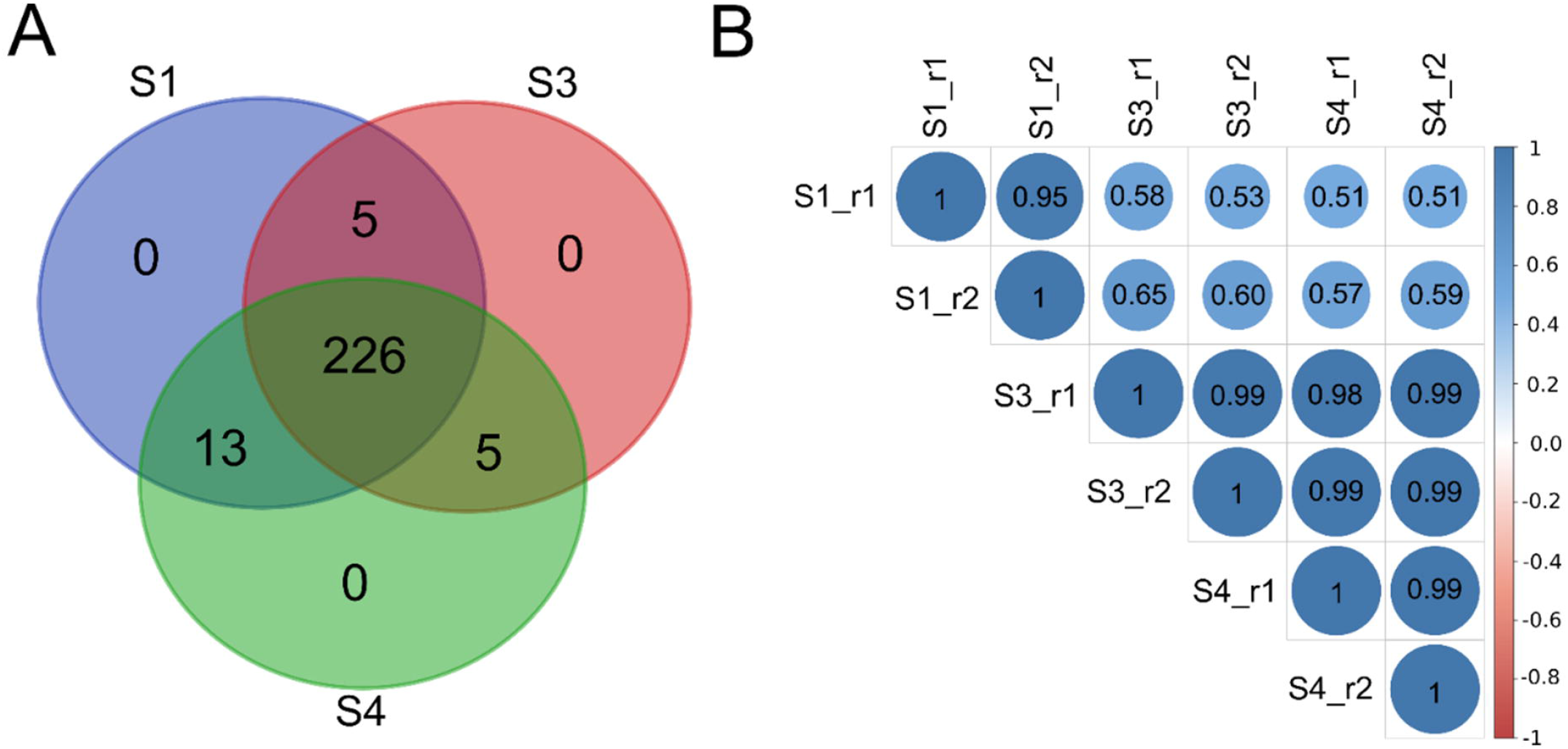
Distribution and correlated expression of genes encoding DOXP/MEP, carotenoid and bixin pathway enzymes between the different seed developmental stages in *B. orellana*. (**A**) Venn diagram of shared genes between the different seed developmental stages (S1, S3, and S4). (**B**) Pearson correlation analysis of gene expression (FDR < 0.05) across all samples.

A total of 50 genes encoding enzymes of the DOXP/MEP, carotenoid, and bixin pathways were identified as DEGs (FDR < 0.05) between all the seed developmental stages (Figs. S3, S4, and S5). Hierarchical clustering analysis of transcriptional profiles showed that half of these DEGs were upregulated during the S1 stage and gradually downregulated during the S3 and S4 stages, whereas the other half showed an opposite behavior (Table S4; Fig. 4). The former included DEGs encoding 19 MTs, three ALDHs, DXS2b, PSY1, and LCY-β1. Of these, 17 MTs (five COMTs, three SABATH-1, two SABATH-12, two CCoAOMT7, MT2×1, PMT28, SABATH-9, SABATH-10, and SAM-dependent MT), all ALDH (two ALDH3F1 and ALDH2C4), DOXP/MEP and carotenoid enzymes showed plastidial localization (Table S4; Fig. 4). The opposite group included DEGs encoding seven CCD4s, five ALDHs, three MTs, DXS2a, DXR, MDS, HDS, IDI, PSY2, ZDS, and LCY-β2. Of these, four ALDHs (three ALDH3I1 and ALDH3F1), three MTs (JAMT, SABATH-3, and SABATH-4), all CCD4s, DOXP/MEP, and carotenoid pathway enzymes exhibited plastidial localization (Table S4; Fig. 4). In addition, two genes (*DXS2a* and *DXS2b*) were upregulated only at the S4 stage (Table S4; Fig. 4).

**Fig. 4.**
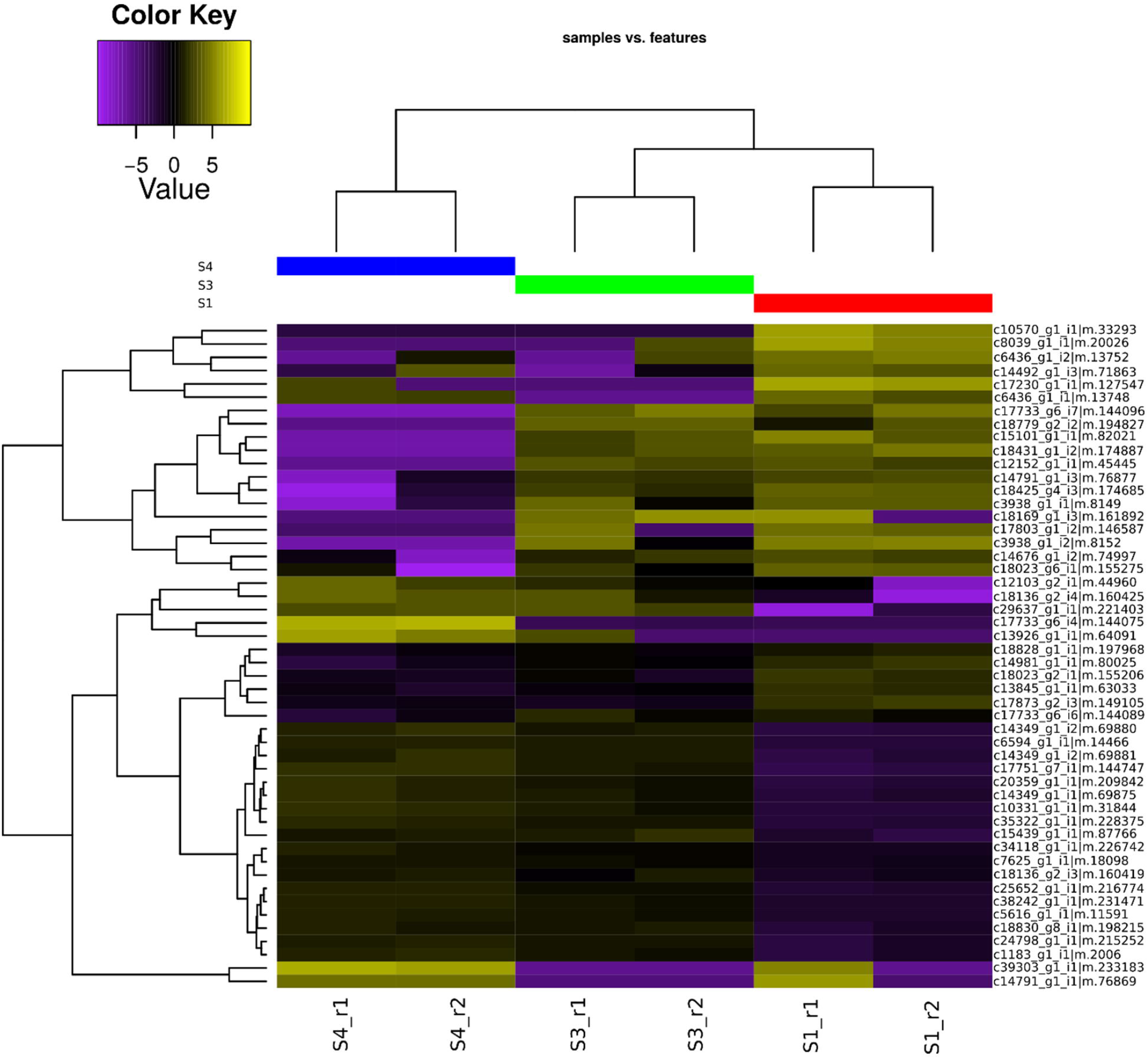
Heatmap of differentially expressed genes (DEGs) encoding DOXP/MEP, carotenoid and bixin pathway enzymes in *B. orellana* at different stages of seed development, with hierarchical clustering in both directions. Color intensity indicates expression strength, yellow corresponds to upregulated genes, and purple denotes downregulated genes (FDR< 0.05). S1, S3, and S4 denote seeds at the first, third, and fourth stages of development, respectively.

### qRT-PCR expression analysis of candidate genes in different organs

Five DEGs identified in our RNA-Seq libraries (*DXS2a, PSY1, ZDS, CCD4-4*, and *SABATH-4*) were selected for qRT-PCR analysis of their expression profiles in different organs of *B. orellana*. Significant (*P* ≤ 0.05) positive correlations (Pearson’s coefficients of 0.75-0.99) were found between the RNA-Seq and qRT-PCR data, confirming the reliability of our transcriptomic data (Fig. S6). *DXS2a, ZDS, CCD4-4*, and *SABATH-4* were preferentially expressed in seeds, and their levels increased throughout seed maturation, peaking at the S4 stage (Fig. 5). In contrast, *PSY1* showed a low but preferential expression in leaves and seeds at the S1 stage, followed by strong downregulation at the S2, S3, and S4 stages (Fig. 5).

**Fig. 5.**
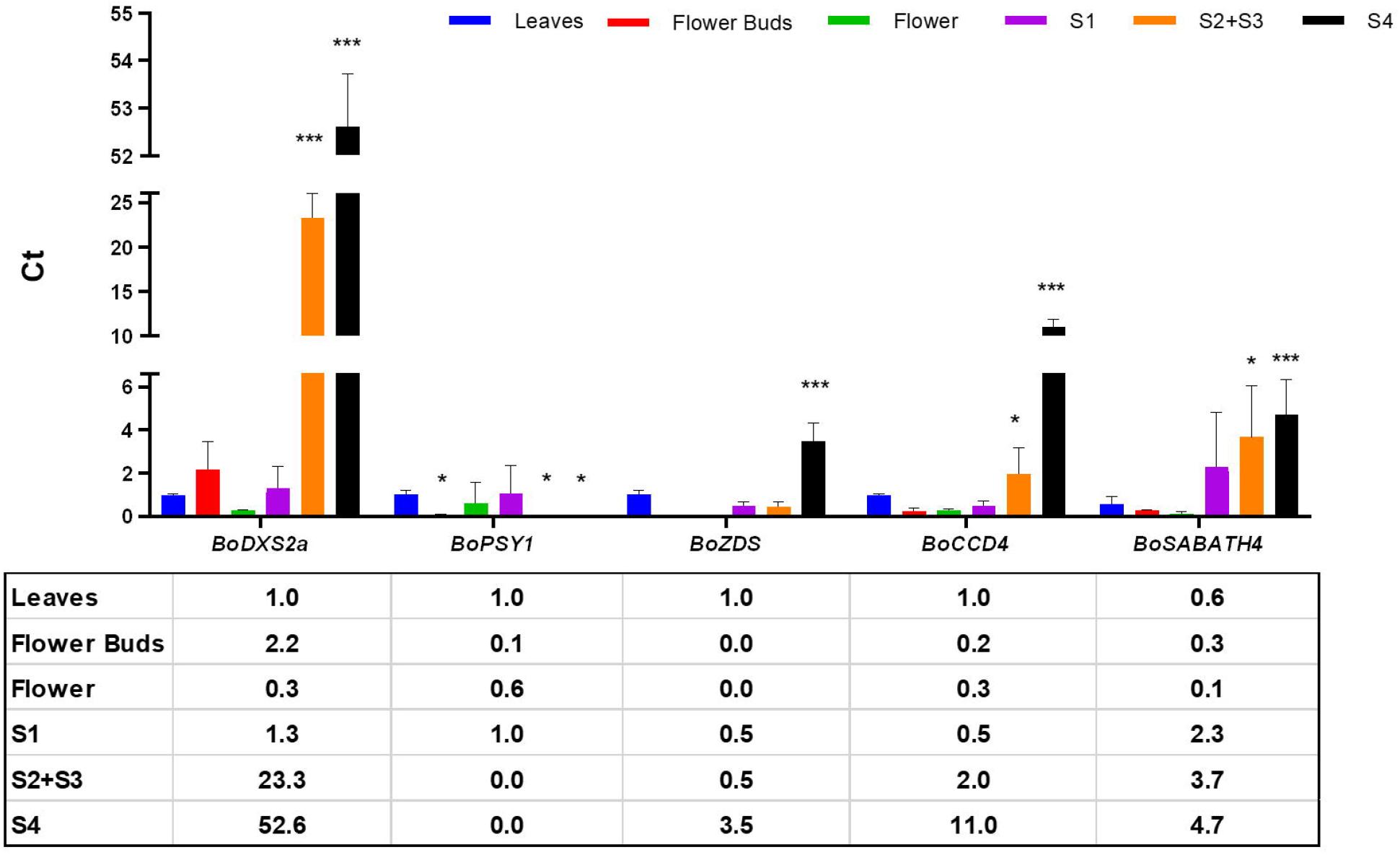
qRT-PCR expression analysis of selected DEGs in different organs of *B. orellana*. Values were normalized to the *RPL38* endogenous control and represent the mean ± SD of three independent biological replicates. \**P* ≤ 0.05 and \*\*\**P* ≤ 0.001 for the leaf (calibrator) vs. different organs according to Student’s *t*-test. S1, S2+S3, and S4, seeds at the first, second plus third, and fourth stages of development.

## Discussion

The results of the present study show that bixin production occurs not only in mature seeds of *B. orellana*, as widely reported (Mercadante and Pfander 1998; Giuliano et al. 2003), but also in all the other organs analyzed (Fig. 1), thus corroborating previous works (Rodríguez-Ávila et al. 2011; Rivera-Madrid et al. 2013). However, only basal levels of bixin were found in mature leaves, flower buds, flowers, and S1-stage seeds; whereas a continuous and massive accumulation of the pigment was observed from the S2 to the S4 stage of seed development, eventually stabilizing at a high level at the mature stage (Fig. 1). Such differences can be attributed to specific CSCs in the aril of *B. orellana* seeds. During development, they differentiate from parenchymatic cells containing starch granules to larger and more numerous cells enabling increased carotenoid accumulation (Louro and Santiago 2016; Carballo-Uicab et al. 2019). Accumulation of elevated levels of bixin at the S5 stage reinforces the idea that mature aril cells lose their biosynthetic functions and serve only for pigment storage (Rodríguez-Ávila et al. 2011).

Our anatomical analysis (Fig. 2) corroborates previous evidences that the massive accumulation of bixin occurs specifically in CSCs in the aril (Louro and Santiago 2016; Carballo-Uicab et al. 2019). These specialized aril cells began to develop asynchronously at the S2 stage, increased in size after fusion of the small vacuoles into a single central vacuole at the S3 stage, further expanded as their large central vacuole became fully filled with bixin to occupy the region of the mesotesta at the S4 stage, and finally matured to the S5 stage with their central vacuole completely filled with bixin (Fig. 2). Thus, the continuous and massive accumulation of bixin observed from the S2 stage onward is a developmentally regulated process that requires the differentiation and development of specialized CSCs capable of synthesizing and storing vast amounts of the pigment. Ultrastructural analyses have shown that during the initial stage of CSC development, in which they display small vacuoles sparsely distributed throughout the cytoplasm, carotenoids accumulate in chromoplasts in the form of electron-dense plastoglobules and electron-dense globules of variable size that surround the plastids (Louro and Santiago 2016). In the next stage, when the small vacuoles fuse to form a single central vacuole, electron-dense lipid bodies are observed in the cytoplasm adjacent to the plastids and tonoplast. In subsequent stages, the CSCs increase in size, displaying a peripheral cytoplasm rich in chromoplasts with numerous plastoglobules in their stroma, as well as large electron-dense lipid bodies resulting from the fusion of the small lipid bodies and electron-lucent vesicles next to the plastids and tonoplast. Additionally, vacuoles become filled with electron-dense globules and electron-lucent vesicles; the former are oil-soluble and contain bixin, whereas the latter are water-soluble and contain norbixin (Louro and Santiago 2016).

In the present study, high-throughput RNA-Seq coupled with chemical and anatomical data from different *B. orellana* seed developmental stages revealed more precisely and comprehensively the key molecular events and candidate genes associated with the developmentally programmed changes in CSCs leading to enhanced bixin synthesis and accumulation. A total of 73,381 protein-encoding contigs were obtained from the transcriptome assembly (Table S3) of seeds at three developmental stages. Their comparison to plant protein databases and functional annotation using the NR, InterPro, and GO databases (Fig. S1) revealed a high degree of similarity with the transcriptome from related plant species, such as *T. cacao* and *Gossypium* spp., as observed previously (Cárdenas-Conejo et al. 2015). Overall, 312 genes encoding DOXP/MEP, carotenoid, and bixin pathway enzymes were identified here, which is more than the 74 genes identified in a previous RNA-Seq study (Cárdenas-Conejo et al. 2015). Similarly, transcripts corresponding to the cDNA sequences previously proposed to be involved in bixin biosynthesis (Bouvier et al. 2003a) were not identified in the present study, suggesting possible mRNA contamination in the earlier work.

Most (90.8%) candidate genes considered for differential expression analysis were present at all seed developmental stages analyzed (Fig. 3A), implying that the DOXP/MEP, carotenoid, and bixin biosynthetic pathways are active in the initial S1 stage (Fig. 1). However, their expression pattern varied over time (Fig. 3B), with 50 of them identified as DEGs between the different seed stages (Table S4; Figs. S3, S4, and S5). These DEGs likely control metabolic flux into carotenoid and bixin pathways during seed development. The subset comprising *DXS2b, PSY1, LCY-β1*, three *ALDH*, and 19 *MT* genes upregulated at the S1 stage and gradually downregulated at the S3 and S4 stages (Table S4; Fig. 4) can be associated with basal levels of bixin found in S1-stage seeds and growing pigment content in S3-stage seeds (Fig. 1). The subset comprising *DXS2a, DXR, MDS, HDS, IDI, PSY2, ZDS, LCY-β2*, seven *CCD4*, five *ALDH*, and three *MT* genes downregulated at the S1 stage and strongly upregulated at the S3 and S4 stages, as well as the two *DXSs* upregulated only at the S4 stage (Table S4; Fig. 4), can be associated with the massive accumulation of bixin found in S3- and S4-stage seeds (Fig. 1). The same subset can be linked also to the reported presence of minor C_40_-carotenoids, such as phytoene, phytofluene, zeta-carotene, neurosporene, β-carotene, cryptoxanthin, and zeaxanthin (Tirimanna 1981; Mercadante and Pfander 1998). Such associations are further supported by the consistent expression patterns of selected genes in different organs (Fig. S6; Fig. 5), as well as by the previous *B. orellana* seed transcriptome (Jako et al. 2002; Soares et al. 2011; Cárdenas-Conejo et al. 2015) and functional (Carballo-Uicab et al. 2019) studies. Except for 11 MTs (five COMTs, two CCoAOMT7, two PMTs, MT2×1, and SAM-dependent MT) identified in the present study, all other DEGs encoding enzymes involved in the DOXP/MEP, carotenoid, and bixin biosynthesis pathways had been identified also in previous *B. orellana* seed transcriptome studies (Jako et al. 2002; Soares et al. 2011; Cárdenas-Conejo et al. 2015), reinforcing their association with bixin biosynthesis. Functional characterization of *B. orellana* CCD4-3 and CCD1-1 has shown that both exhibit tissue-specific expression in the CSCs of seed aril and in vitro lycopene cleavage activity (Carballo-Uicab et al. 2019). Furthermore, similar functions have been described for homologous sequences from other plant species. For example, DXS2, but not DXS1 or DXS3, appear involved in carotenoids formation in non-photosynthetic tissues such as seeds (Peng et al. 2013; Saladié et al. 2014); LCY-β1 plays such role in chloroplast-containing tissues while LCY-β2 only in chromoplast-containing tissues (Ronen et al. 2000; Galpaz et al. 2006). CCD4, but not CCD1 or other CCDs, participate in the formation of unique apocarotenoid pigments found in flower and fruit tissues (Ohmiya 2009; Hou et al. 2016). ALDH2C4 contributes to terpenoid metabolism (Nair et al. 2004; Xie et al. 2018), while ALDH3I1 to the oxidation of crocetin dialdehyde to crocetin in saffron (Tola et al. 2021). COMTs participate in the phenylpropanoid and volatile floral scent-determining metabolic pathways (Wu et al. 2003; Han et al. 2007); whereas SABATHs are involved in the methylation of carboxyl groups and nitrogen moieties of a variety of low-molecular weight metabolites (D’Auria et al. 2003).

The subcellular compartmentalization of the bixin pathway remains an open question. In the seeds of *B. orellana*, carotenoids are synthesized within chromoplasts, with the final products stored in the vacuole (Bouvier et al. 2003b; Giuliano et al. 2003; Louro and Santiago 2016). In contrast to the CCD4 clade enzymes, which are localized in plastoglobules within plastids, the enzymes catalyzing the last two steps in the *B. orellana* bixin pathway, bixin aldehyde dehydrogenase and norbixin methyltransferase, do not carry recognizable plastid transit peptides (Bouvier et al. 2003a), suggesting that these steps could occur in the cytoplasm (Giuliano et al. 2003). However, we report here a predicted plastidial localization for seven ALDHs (three ALDH3F1, three ALDH3I1, and ALDH2C4) and 20 MTs (five COMTs, three SABATH-1, two SABATH-12, two CCoAOMT7, JAMT, MT2×1, PMT28, SABATH-3, SABATH-4, SABATH-9, SABATH-10, and SAM-dependent MT) (Table S4), in addition to all CCD4s. These findings suggest that the entire bixin pathway may take place inside plastids, with subsequent transfer of the pigments to the vacuole via large lipid bodies and vesicles (Louro and Santiago 2016). A model of crocin transport from the chromoplast to the vacuole in saffron stigmas was recently proposed (Gómez-Gómez et al. 2017). This model is based on the direct transfer of large globules containing crocins, named crocinoplasts and originating from the fusion of many vesicle-like structures inside the plastid, from the polarized end of the chromoplast to the vacuole, in a process involving the action of vesicle fusion-mediating proteins and tonoplast receptors (Gómez-Gómez et al. 2017). Whether a similar form of apocarotenoid transport from the chromoplast to the vacuole could operate for bixin remains to be confirmed.

Integration of our present chemical, anatomical, and transcriptomic data with existing literature allows us to propose a refined model of the key cellular and molecular events contributing to bixin synthesis and accumulation (Fig. 6). It is based on the developmentally regulated processes of CSC differentiation and development, as well as the coordinated expression of DOXP/MEP, carotenoid, and bixin biosynthesis genes observed throughout the different stages of seed development, and culminating in pigment accumulation in the vacuole. According to this model, in the first stage of seed development, CSC precursor parenchyma cells contain amyloplasts with few and small plastoglobules (Louro and Santiago 2016), with the DOXP/MEP, carotenoid, and bixin biosynthetic pathways operating at a limited rate and driven by *DXS2b, PSY1, LCY-β1, ALDH2C4, ALDH3F1, COMT*s, *CCoAOMT7, MT2×1, PMT28, SABATH-1/9/10/12*, and *SAM-dependent MT* (Table S4; Fig. 4). From this stage onward, CSCs begin to differentiate and develop (Fig. 2), with the former group of genes being progressively turned off, while *DXS2a, DXR, MDS, HDS, IDI, PSY2, ZDS, LCY-β2, CCD4*s, *ALDH3F1, ALDH3I1, JAMT*, and *SABATH-3/4* are progressively turned on (Table S4; Fig. 4). This leads to the massive accumulation of carotenoids in chromoplasts in the form of plastoglobules (Louro and Santiago 2016), which gradually fuse together to generate large globules and vesicles that are subsequently transferred inside the vacuole (Louro and Santiago 2016; Gómez-Gómez et al. 2017). Our model does not exclude the possibility that the last two steps in the bixin pathway could also take place in the cytoplasm, in a process that might involve the transient association of cytosolic CCDs, ALDHs and MTs, with the outer envelope of the chromoplasts to access bixin aldehyde.

**Fig. 6.**
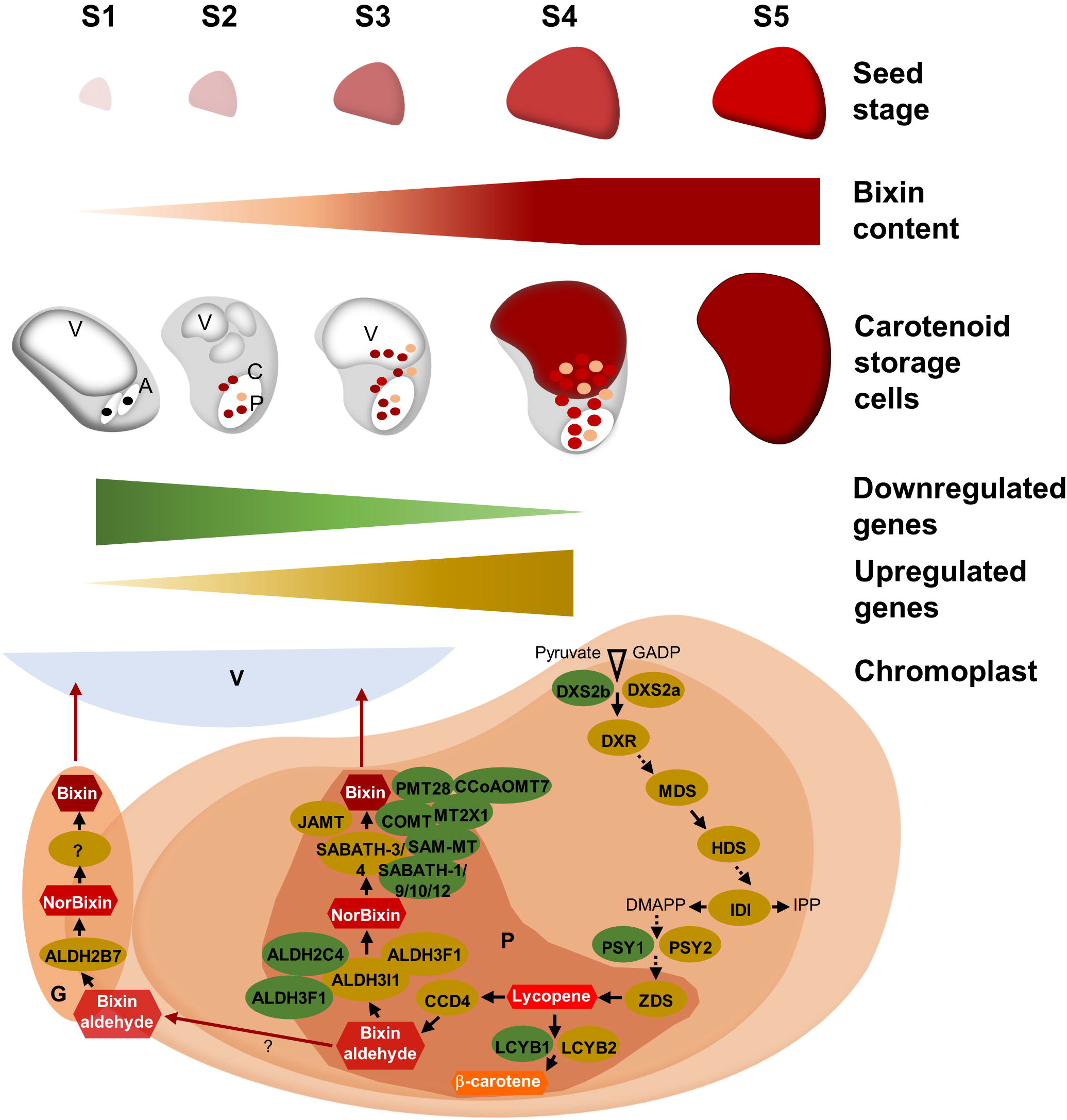
Proposed model of bixin synthesis and accumulation in seeds of *B. orellana*. At the first stage of development, the seed aril comprises only CSC precursor parenchyma cells capable of limited pigment synthesis and accumulation associated with the few and small plastoglobules (P) inside amyloplasts (A). Metabolic flux into carotenoid and bixin biosynthesis is controlled by the transcriptional activation of genes marked in green. From this stage onward, CSCs begin to differentiate and develop, with the former group of genes progressively turned-off and the genes marked in yellow progressively turned-on. This leads to the massive accumulation of carotenoids in plastoglobules inside chromoplasts (C) and also in cytoplasmic vesicles and lipid globules (G) that surround plastids and the tonoplast. The latter gradually fuse together to generate large globules and vesicles that are subsequently transferred inside the vacuole (V). (Apo-)carotenoids are depicted inside hexagons. Black solid-line arrows indicate a single enzymatic step and dashed-line arrows indicate more than one enzymatic step. ALDH, aldehyde dehydrogenase; BAM-MT, benzoic acid carboxyl methyltransferase; CCD, carotenoid cleavage dioxygenase; CCoAOMT, caffeoyl coenzyme A ester O-methyltransferase; COMT, caffeic acid 3-O-methyltransferase; DMAPP, dimethylallyl pyrophosphate; DXR, 1-deoxy-D-xylulose-5-phosphate reductoisomerase; DXS, 1-deoxy-D-xylulose-5-phosphate synthase; GADP, glyceraldehyde-3-phosphate; HDS, 4-hydroxy-3-methylbut-2-en-1-yl diphosphate synthase; IDI, isopentenyl diphosphate isomerase; IPP, isopentenyl pyrophosphate; JAMT, jasmonic acid carboxyl methyltransferase; LCYB, lycopene beta-cyclase; MDS, 2-C-methyl-D-erythritol 2,4-cyclodiphosphate synthase; MT2×1, methyltransferase 2 isoform X1; PMT, putrescine N-methyltransferase; PSY, phytoene synthase; SABATH, SAM-MT/BAM-MT/Theobromine synthase methyltransferase; SAM-MT, salicylic acid carboxyl methyltransferase; ZDS, zeta-carotene desaturase.

## Conclusion

The present study advances our understanding of the cellular and molecular events and candidate genes associated with bixin synthesis and accumulation during seed development in *B. orellana*. Additionally, it opens new questions regarding the processes and genes/proteins involved in CSC differentiation and bixin transport to the vacuole. From a biotechnological point of view, enhancing the metabolic sink strength for bixin in chromoplasts and vacuoles, particularly through induction of CSC formation along with overexpression of the genes identified in the present study, could significantly enhance bixin content in annatto or help engineer the pathway in other organisms.

## Supplementary information

The online version contains supplementary material available at

## Acknowledgments

We gratefully acknowledge the material and technical support provided by Carlos Priminho Pirovani (Universidade Estadual de Santa Cruz, Ilhéus, Bahia, Brazil). We would like to thank Editage (www.editage.com) for English language editing.

## Author contribution statement

VSM conducted the experiments. VSM, VLFS, VCS, AOS, KTSN, and MRS analyzed the data. VSM drafted the manuscript. VLFS, PVZCG, TNHR, CAEL, DVF, WCO, and MGCC supported the project and designed the experiments. VLFS, WCO, and MGCC revised the manuscript. All authors read and approved the manuscript.

## Supplementary data

**Table S1** Genes and primers used for qRT-PCR reactions.

**Table S2** Sequencing results for *B. orellana* seeds at three developmental stages (S1, S3, and S4), obtained from two biological replicates (r1 and r2), and generated using the Illumina HiSeq 2500 platform, with 150-bp paired-end reads. G, giga; Q, quality.

**Table S3** Assembly statistics of the *B. orellana* seed transcriptome.

**Table S4** Differentially expressed genes related to the DOXP/MEP, carotenoid, and bixin biosynthesis pathways.

**Fig. S1** Functional annotation of the *B. orellana* seed transcriptome. (**A**) Top-hits species distribution. (**B**) Number of sequences annotated against the NR, InterPro, and GO databases.

**Fig. S2** KEGG pathways showing transcripts/enzymes identified in the DOXP/MEP and carotenoid pathways. Enzymes with Enzyme Commission number highlighted in different color were identified in the present study.

**Fig. S3** Analysis of differentially expressed genes (FDR < 0.05) between S1 and S3 stages of seed development. Red dots indicate differentially expressed contigs, whereas black dots indicate nonsignificant genes.

**Fig. S4** Analysis of differentially expressed genes (FDR < 0.05) between S1 and S4 stages of seed development. Red dots indicate differentially expressed contigs whereas black dots indicate nonsignificant genes.

**Fig. S5** Analysis of differentially expressed genes (FDR < 0.05) between S3 and S4 stages of seed development. Red dots indicate differentially expressed contigs whereas black dots indicate nonsignificant genes.

**Fig. S6** Pearson’s correlations (R) between the levels of gene expression measured by RNA-Seq and qRT-PCR.

