## Supplemental File for "Dynamics of annatto pigment synthesis and accumulation revealed by integrated chemical, anatomical, and RNA-Seq analyses"

##### **Supplementary data (Moreira et al.)**

##### **Table S1** Genes and primers used for qRT-PCR reactions

| **Gene** | **Sequence 5´- 3´** | | **GenBank accession**  **no.** | **Tm**  **(ºC)** | **Ampli-con size**  **(pb)** | **Efficiency (E)** |
| --- | --- | --- | --- | --- | --- | --- |
| ***CCD4-4*** | *F-* *GCGAAGTTGGAAGCAGATTCTT*  *R-* *ACATGGAATCCCCACTATTTTCA* | | MW885484 | 55.8  54.3 | 150 | 83.98 |
| ***PSY1*** | *F-* *CGATGAGGCAGAGAAAGGAG*  *R-* *TGGCGATACAACAACAAGGA* | | MW885381 | 55  54.4 | 81 | 83.81 |
| ***DXS2a*** | *F- CAGGCATGGCTTTCACTT*  *R- CGGCAAAGGTAGATGAGTATG* | | MW885565 | 53.3  53 | 150 | 84.15 |
| ***ZDS*** | *F-TCCTCTCCATGGGATTCAAG*  *R-GACTGAAGGCAAGAGCCAAA* | | MW885557 | 53.8  55.5 | 89 | 82.79 |
| ***SABATH4*** | *F-TGCTCCTCAGGACCTACTGC*  *R-GCAATGCTGTTGTTGGCATA* | | MW885576 | 58.7  54.3 | 93 | 83.93 |
| ***RPL38*** | *F-ACTTCCTTCTCACCGCCAGA*  *R-CAGCGGACCTTGAACTTGAC* | MW736056 | | 58.3  57.2 | 85 | 83.77 |

##### **Table S****2** Sequencing results for *B. orellana* seeds at three developmental stages (S1, S3, and S4), obtained from two biological replicates (r1 and r2), and generated using the Illumina HiSeq 2500 platform, with 150-bp paired-end reads. G, giga; Q, quality

| **Sample** | **Raw reads** | **Clean reads** | **Raw base**  **(G)** | **Clean base**  **(G)** | **Effective rate**  **(%)** | **Q20**  **(%)** | **Q30**  **(%)** | **GC**  **(%)** |
| --- | --- | --- | --- | --- | --- | --- | --- | --- |
| S1_r1 | 2,308,303 | 1,249,771 | 0.69 | 0.37 | 54.14 | 97.27 | 92.92 | 45.12 |
| S1_r2 | 2,291,560 | 1,158,931 | 0.69 | 0.35 | 50.57 | 97.41 | 93.21 | 45.12 |
| S3_r1 | 2,222,017 | 1,281,908 | 0.67 | 0.38 | 57.69 | 96.75 | 91.95 | 45.39 |
| S3_r2 | 2,327,846 | 1,260,156 | 0.70 | 0.38 | 54.13 | 97.30 | 93.06 | 45.32 |
| S4_r1 | 2,478,894 | 1,565,215 | 0.74 | 0.47 | 63.14 | 97.35 | 93.22 | 45.45 |
| S4_r2 | 1,813,470 | 1,056,613 | 0.54 | 0.32 | 58.26 | 97.37 | 93.23 | 45.15 |
| TOTAL | 13,442,090 | 7,572,594 | 4.03 | 2.27 | - | - | - | - |

##### **Table S3** Assembly statistics of the *B. orellana* seed transcriptome

| Transcriptome *de novo* data | All contigs | Longest isoform of the ´gene´. |
| --- | --- | --- |
| Total number of bases assembled (bp) | 52,471,683 | 22,234,446 |
| **Total number of assembled contigs** | 79,650 | 46,115 |
| GC Percentage | 42.10 | 42.10 |
| *Contig* N10 | 3,500 | 3,537 |
| *Contig* N20 | 2,736 | 2,735 |
| *Contig* N30 | 2,266 | 2,232 |
| *Contig* N40 | 1,911 | 1,855 |
| *Contig* N50 | 1,603 | 1,515 |
| *Contig* average (bp) | 994.91 | 827.65 |
| **Total number of ORFs** | 73,381 | 26,908 |
| GC Percentage | 44 | 44 |
| *ORF* N10 | 2,919 | 3,120 |
| *ORF* N20 | 2,175 | 2,346 |
| *ORF* N30 | 1,752 | 1,899 |
| *ORF* N40 | 1,446 | 1,575 |
| *ORF* N50 | 1,191 | 1,335 |
| *ORF* average (bp) | 715.06 | 826.31 |

**Table S4** Differentially expressed genes related to the DOXP/MEP, carotenoid, and bixin biosynthesis pathways

| **Gene ID** | **GenBank accession no.** | **Name** | **Ontology** | **Predicted subcellular localization** |
| --- | --- | --- | --- | --- |
| ***Up- to down-regulated genes from S1- to S4-stage*** | | | | |
| *DOXP/MEP pathway* | | | | |
| >c14791_g1_i3\|m.76877 | MW885368 | 1-deoxy-D-xylulose-5-phosphate synthase 2b | GO:0008661 GO:0016114 | Plastid |
| *Carotenoid pathway* | | | | |
| >c14981_g1_i1\|m.80025 | MW885381 | Phytoene synthase 1 | GO:0010287 GO:0004310 GO:0016767 GO:0051996 GO:0006696 GO:0016117 | Plastid |
| >c14676_g1_i2\|m.74997 | MW885359 | Lycopene beta-cyclase 1 | GO:0009507 GO:0016491 GO:0055114 | Plastid |
| *Bixin pathway* | | | | |
| >c13845_g1_i1\|m.63033 | MW885330 | Aldehyde dehydrogenase family 2 member C4 | GO:0001758 GO:0055114 | Plastid |
| >c17733_g6_i6\|m.144089 | MW885465 | Aldehyde dehydrogenase family 3 member F1 | GO:0004028 GO:0004029 GO:0004030 GO:0006081 GO:0055114 | Plastid |
| >c17733_g6_i7\|m.144096 | MW885466 | Aldehyde dehydrogenase family 3 member F1 | GO:0004028 GO:0004029 GO:0004030 GO:0006081 GO:0055114 | Plastid |
| >c6436_g1_i1\|m.13748 | MW885574 | Caffeic acid 3-O-methyltransferase 1 | GO:0005829 GO:0008171 GO:0008757 GO:0046983 GO:0019438 GO:0032259 | Plastid |
| >c6436_g1_i2\|m.13752 | MW885575 | Caffeic acid 3-O-methyltransferase-like | GO:0005829 GO:0046983 GO:0047763 GO:0032259 | Plastid |
| >c18023_g2_i1\|m.155206 | MW885478 | Caffeic acid 3-O-methyltransferase-like | GO:0008171 GO:0046983 GO:0032259 | Plastid |
| >c18023_g6_i1\|m.155275 | MW885479 | Caffeic acid 3-O-methyltransferase-like | GO:0008171 GO:0046983 GO:0032259 | Plastid |
| >c18425_g4_i3\|m.174685 | MW885494 | Caffeic acid 3-O-methyltransferase-like | GO:0008171 GO:0046983 GO:0032259 | Plastid |
| >c17873_g2_i3\|m.149105 | MW885477 | Caffeoyl coenzyme A ester O-methyltransferase 7 | GO:0005829 GO:0042409 GO:0032259 | Plastid |
| >c18828_g1_i1\|m.197968 | MW885512 | Caffeoyl coenzyme A ester O-methyltransferase 7 | GO:0005829 GO:0042409 GO:0032259 | Plastid |
| >c14492_g1_i3\|m.71863 | MW885343 | Methyltransferase 2 isoform X1 | GO:0003676 GO:0008168 GO:0006139 GO:0032259 | Plastid |
| >c18779_g2_i2\|m.194827 | MW885509 | Methyltransferase PMT7 | GO:0005768 GO:0005774 GO:0005802 GO:0016021 GO:0008757 GO:0032259 | Endoplasm |
| >c17803_g1_i2\|m.146587 | MW885469 | Methyltransferase PMT28 | GO:0005768 GO:0005802 GO:0016021 GO:0008168 GO:0010181 GO:0051536 GO:0032259 GO:0055114 | Plastid |
| >c3938_g1_i1\|m.8149 | MW885566 | SABATH methyltransferase 1 | GO:0008168 GO:0032259 | Plastid |
| >c3938_g1_i2\|m.8152 | MW885567 | SABATH methyltransferase 1 | GO:0008168 GO:0032259 | Plastid |
| >c10570_g1_i1\|m.33293 | MW885280 | SABATH methyltransferase 1 | GO:0005829 GO:0008757 GO:0032259 | Plastid |
| >c8039_g1_i1\|m.20026 | MW885580 | SABATH methyltransferase 2 | GO:0008168 GO:0032259 | Nucleus |
| >c17230_g1_i1\|m.127547 | MW885436 | SABATH methyltransferase 9 | GO:0008168 GO:0032259 | Plastid |
| >c12152_g1_i1\|m.45445 | MW885301 | SABATH methyltransferase 10 | GO:0008168 GO:0032259 | Plastid |
| >c15101_g1_i1\|m.82021 | MW885390 | SABATH methyltransferase 12 | GO:0008168 GO:0032259 | Plastid |
| >c18169_g1_i3\|m.161892 | MW885488 | SABATH methyltransferase 12 | GO:0008168 GO:0032259 | Plastid |
| >c18431_g1_i2\|m.174887 | MW885495 | S-adenosyl-L-methionine-dependent methyltransferase | GO:0005768 GO:0005774 GO:0005802 GO:0008757 GO:0032259 | Plastid |
| ***Down- to up-regulated genes from S1- to S4-stage*** | | | | |
| *DOXP/MEP pathway* | | | | |
| >c20359_g1_i1\|m.209842 | MW885531 | 1-deoxy-D-xylulose-5-phosphate synthase 2a | GO:0008661 GO:0016114 | Plastid |
| >c38242_g1_i1\|m.231471 | MW885564 | 1-deoxy-D-xylulose 5-phosphate reductoisomerase | GO:0005515 GO:0016853 GO:0030145 GO:0030604 GO:0070402 GO:0051484 GO:0055114 | Plastid |
| >c34118_g1_i1\|m.226742 | MW885554 | 2-C-methyl-D-erythritol 2,4-cyclodiphosphate synthase | GO:0009570 GO:0008685 GO:0015995 GO:0016117 | Plastid |
| >c5616_g1_i1\|m.11591 | MW885570 | 4-hydroxy-3-methylbut-2-en-1-yl diphosphate synthase | GO:0009570 GO:0009941 GO:0003725 GO:0005506 GO:0046429 GO:0051539 GO:0009617 GO:0009862 GO:0016114 GO:0055114 | Plastid |
| >c7625_g1_i1\|m.18098 | MW885579 | Isopentenyl diphosphate Delta-isomerase I | GO:0004452 GO:0016787 GO:0008299 | Plastid |
| *Carotenoid pathway* | | | | |
| >c10331_g1_i1\|m.31844 | MW885277 | Phytoene synthase 2 | GO:0009536 GO:0004310 GO:0051996 GO:0006696 | Plastid |
| >c35322_g1_i1\|m.228375 | MW885557 | Zeta-carotene desaturase | GO:0009509 GO:0009941 GO:0016719 GO:0052886 GO:0052887 GO:0016117 GO:0052889 GO:0055114 | Plastid |
| >c18830_g8_i1\|m.198215 | MW885513 | Chromoplast-specific lycopene beta-cyclase 2 | GO:0009509 GO:0016705 GO:0052727 GO:0052728 GO:0016117 GO:0055114 | Plastid |
| *Bixin pathway* | | | | |
| >c25652_g1_i1\|m.216774 | MW885538 | Carotenoid cleavage dioxygenase 4 isoform 1 | GO:0016702 GO:0055114 | Plastid |
| >c17751_g7_i1\|m.144747 | MW885467 | Carotenoid cleavage dioxygenase 4 isoform 2 | GO:0016702 GO:0055114 | Plastid |
| >c24798_g1_i1\|m.215252 | MW885537 | Carotenoid cleavage dioxygenase 4 isoform 3 | GO:0005515 GO:0016702 GO:0055114 | Plastid |
| >c18136_g2_i3\|m.160419 | MW885484 | Carotenoid cleavage dioxygenase 4 isoform 4 | GO:0016702 GO:0055114 | Plastid |
| >c18136_g2_i4\|m.160425 | MW885485 | Carotenoid cleavage dioxygenase 4 isoform 4 | GO:0016702 GO:0055114 | Plastid |
| >c29637_g1_i1\|m.221403 | MW885544 | Carotenoid cleavage dioxygenase 4 isoform 4 | GO:0016702 GO:0055114 | Plastid |
| >c15439_g1_i5\|m.87766 | MW885397 | Carotenoid cleavage dioxygenase 4 isoform 5 | GO:0016702 GO:0055114 | Plastid |
| >c13926_g1_i1\|m.64091 | MW885331 | Aldehyde dehydrogenase family 2 member B7 | GO:0004029 GO:0055114 | Cytoplasm |
| >c17733_g6_i4\|m.144075 | MW885463 | Aldehyde dehydrogenase family 3 member F1 | GO:0004028 GO:0004029 GO:0004030 GO:0006081 GO:0055114 | Plastid |
| >c14349_g1_i1\|m.69875 | MW885337 | Aldehyde dehydrogenase family 3 member I1 | GO:0009941 GO:0004028 GO:0004029 GO:0004030 GO:0033721 GO:0006081 GO:0055114 | Plastid |
| >c14349_g1_i2\|m.69880 | MW885338 | Aldehyde dehydrogenase family 3 member I1 | GO:0009941 GO:0004028 GO:0004029 GO:0004030 GO:0033721 GO:0006081 GO:0055114 | Plastid |
| >c14349_g1_i2\|m.69881 | MW885339 | Aldehyde dehydrogenase family 3 member I1 | GO:0004028 GO:0004029 GO:0004030 GO:0006081 GO:0055114 | Plastid |
| >c12103_g2_i1\|m.44960 | MW885300 | Jasmonate O-methyltransferase-like | GO:0030795 GO:0032259 | Plastid |
| >c1183_g1_i1\|m.2006 | MW885294 | SABATH methyltransferase 3 | GO:0008168 GO:0032259 | Plastid |
| >c6594_g1_i1\|m.14466 | MW885576 | SABATH methyltransferase 4 | GO:0008168 GO:0032259 | Plastid |
| ***Upregulated genes only at S4-stage*** | | | | |
| *DOXP/MEP pathway* | | | | |
| >c39303_g1_i1\|m.233183 | MW885565 | 1-deoxy-D-xylulose-5-phosphate synthase 2a | GO:0009536 GO:0008661 GO:0016114 | Plastid |
| >c14791_g1_i1\|m.76869 | MW885366 | 1-deoxy-D-xylulose-5-phosphate synthase 2b | GO:0008661 GO:0016114 | Plastid |

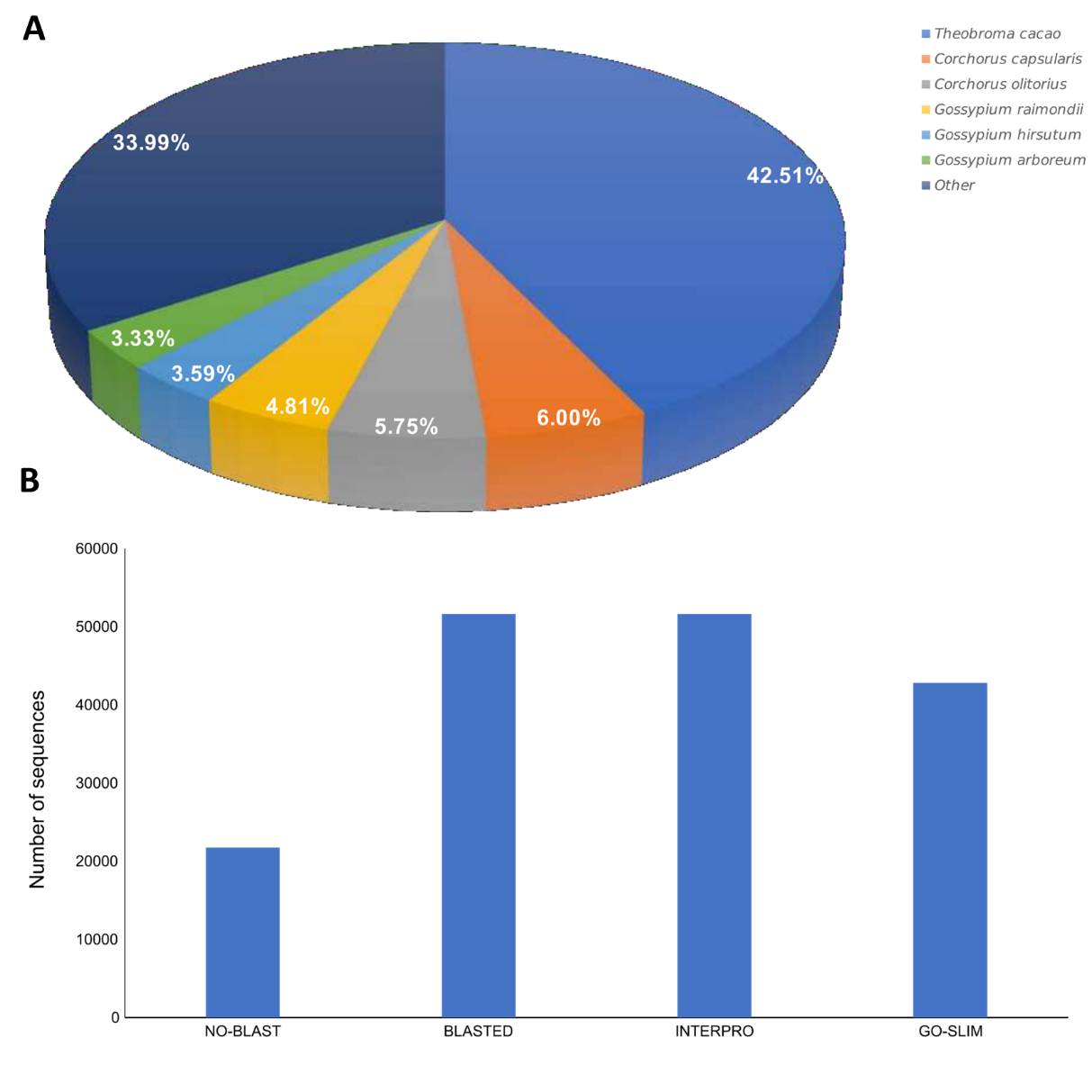

**Fig. S1** Functional annotation of the *B. orellana* seed transcriptome. (**A**) Top-hits species distribution. (**B**) Number of sequences annotated against the NR, InterPro, and GO databases.

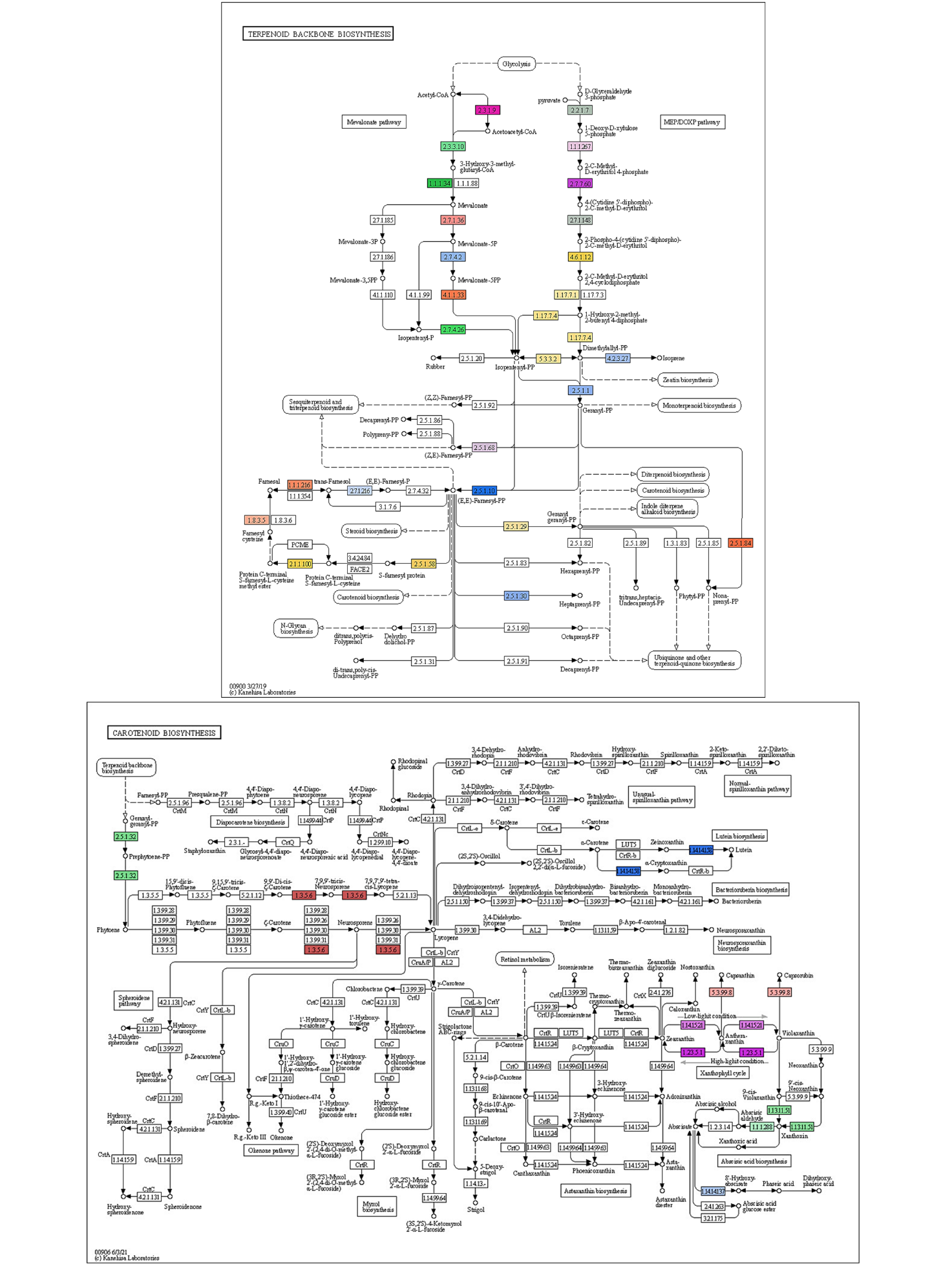
**Fig. S2** KEGG pathways showing transcripts/enzymes identified in the DOXP/MEP and carotenoid pathways. Enzymes with Enzyme Commission number highlighted in different color were identified in the present study.

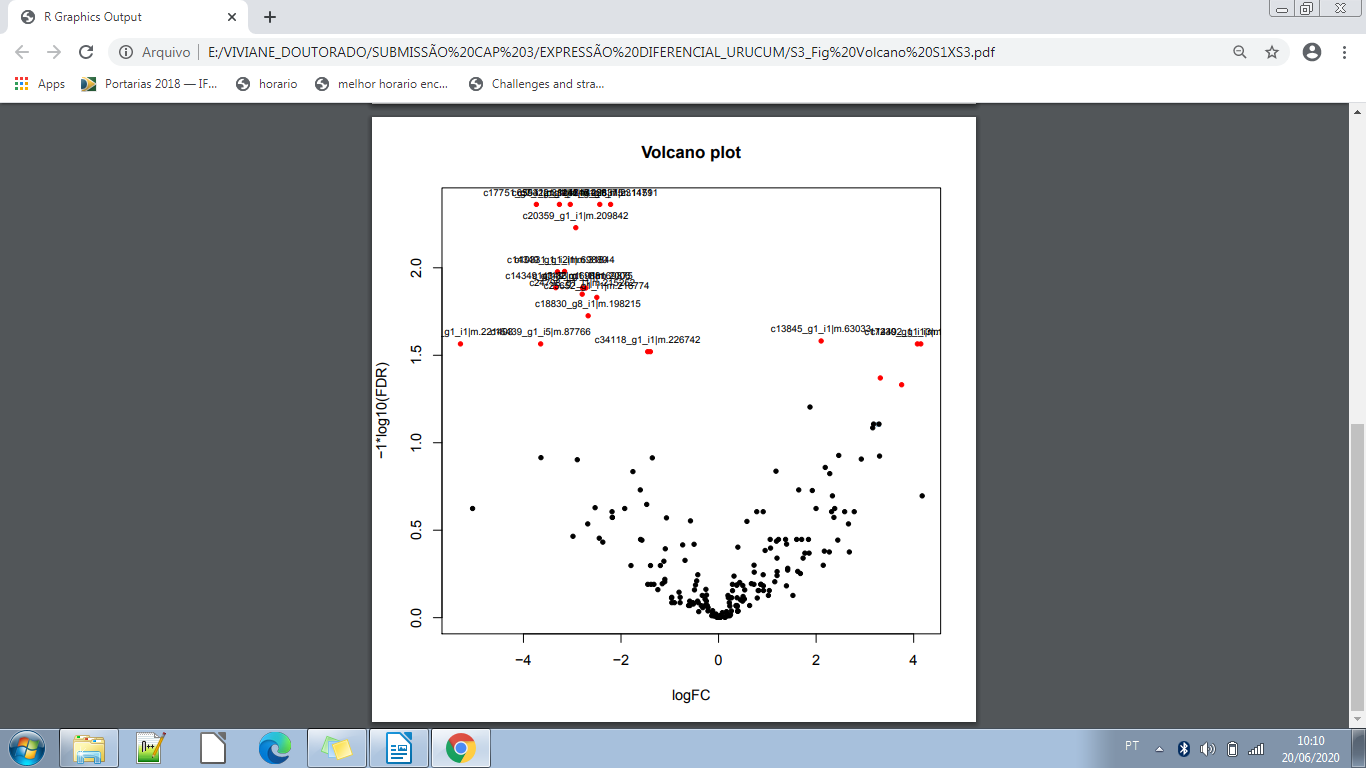

**Fig. S3** Analysis of differentially expressed genes (FDR < 0.05) between S1 and S3 stages of seed development. Red dots indicate differentially expressed contigs, whereas black dots indicate nonsignificant genes.

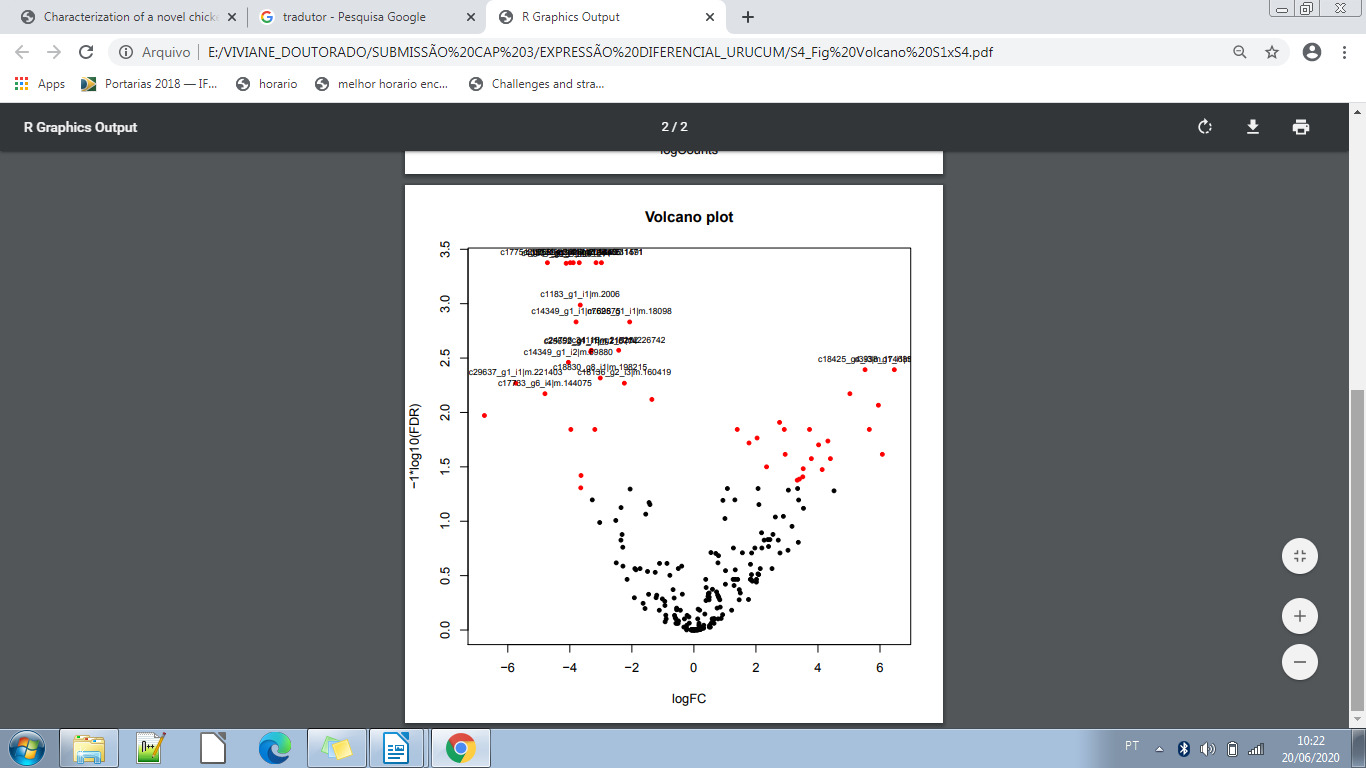

**Fig. S4** Analysis of differentially expressed genes (FDR < 0.05) between S1 and S4 stages of seed development. Red dots indicate differentially expressed contigs whereas black dots indicate nonsignificant genes.

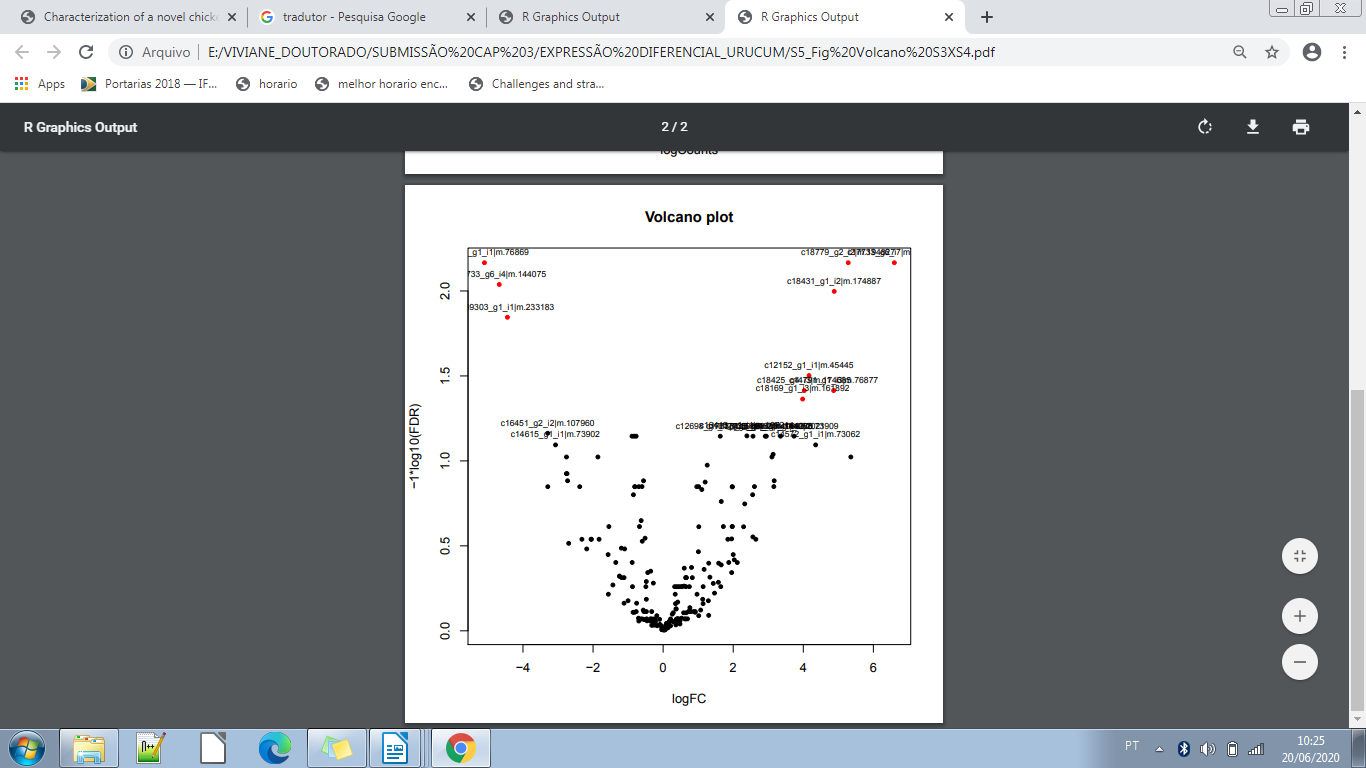

**Fig. S5** Analysis of differentially expressed genes (FDR < 0.05) between S3 and S4 stages of seed development. Red dots indicate differentially expressed contigs whereas black dots indicate nonsignificant genes.

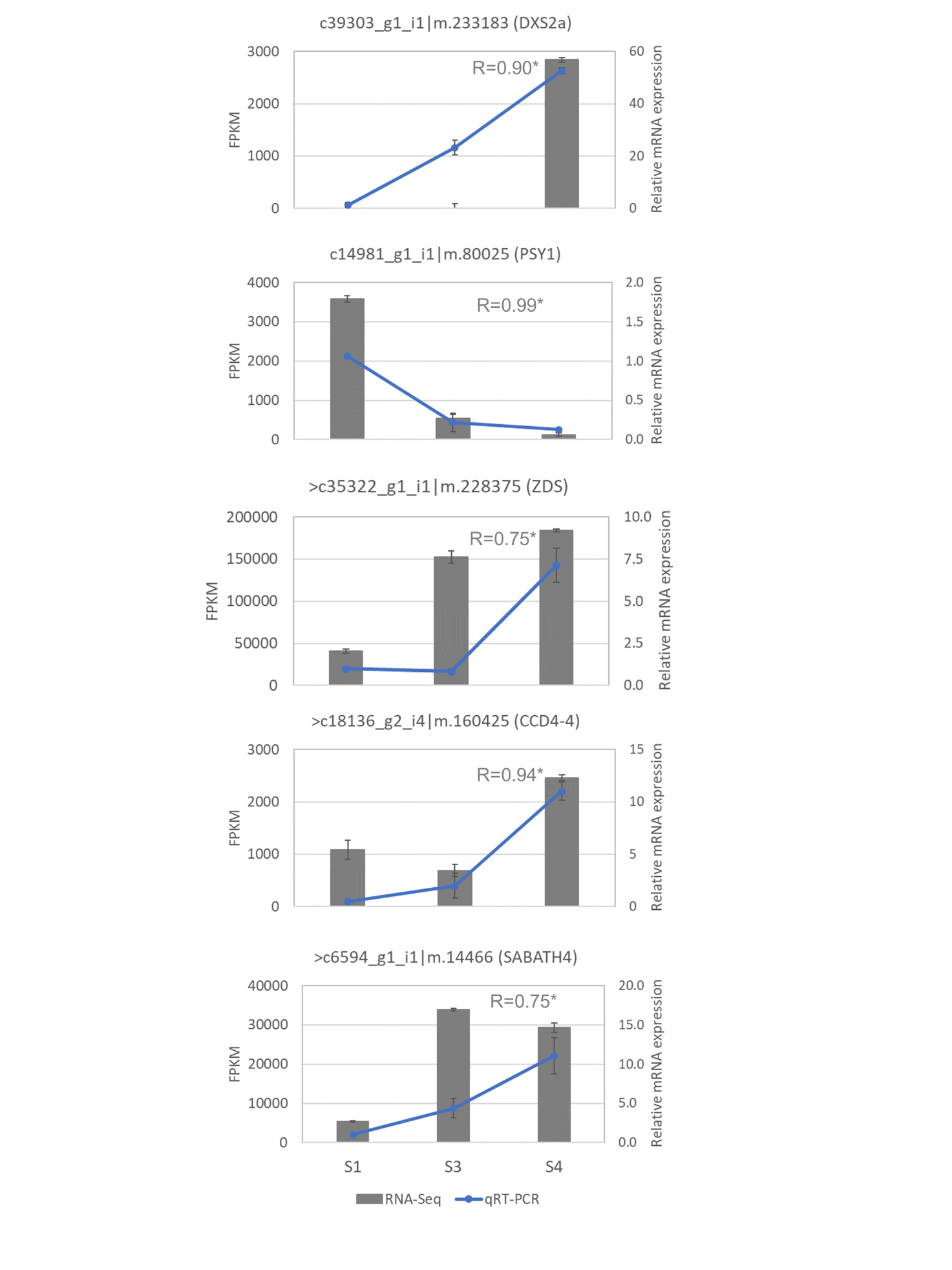

**Fig S6** Pearson’s correlations (R) between the levels of gene expression measured by RNA-Seq and qRT-PCR.
